# An Evolutionary Statistics Toolkit for Simplified Sequence Analysis on Web with Client-Side Processing

**DOI:** 10.1101/2024.08.01.606148

**Authors:** Alper Karagöl, Taner Karagöl

## Abstract

We present the Evolutionary Statistics Toolkit, a user-friendly web-based platform designed for specialized analysis of genetic sequences, which integrates multiple evolutionary statistics. The toolkit focuses on a selection of specialized tools, including Tajima’s D calculator with Site Frequency Spectrum (SFS), Shannon’s Entropy (H), alignment re-formatting, HGSV to FASTA conversion, pair-wise frequency analysis, FASTA to SEQRES, RNA 2D structure alignment, Kyte-Doolittle hydrophilicity plot tool, Chou-Fasman tool, and kurtosis coefficient calculator. Tajima’s D is calculated using the reference formula: D = (π - θ_W_) / sqrt(V_D_), where π corresponds to the average number of differences, θ_W_ is Watterson’s estimator of θ, and V_D_ is the variance of π - θ_W_. Shannon’s Entropy is defined as H = −∑ p_i_ * log_2_(p_i_), where p_i_ is the probability of occurrence of each unique character (nucleotide or amino acid) in the sequence. The toolkit facilitates streamlined workflows for early researchers in evolutionary biology, genomics, and related fields. With comparing with existing codes, we propose it also emerges as an educational interactive website for beginners in evolutionary statistics. The source code for each tool in the toolkit is available through GitHub links provided on the website. This open-source approach allows users to inspect the code, suggest improvements, or further adapt the tools for their specific usage and research needs. This article describes the functionalities, and validation of each tool within the platform, along with comparison with accessible existing statistical utilities. The toolkit is freely accessible on: https://www.alperkaragol.com/toolkit

## Introduction

Understanding the evolutionary dynamics of genetic sequences is fundamental in evolutionary biology and genomics [1]. Contrary to their fundamental roles, traditional tools for sequence analysis often require extensive computational expertise and composed of various independent fragmented workflows [2,3]. To address these challenges, we developed the “Evolutionary Statistics Toolkit,” an integrated platform that simplifies the analysis while providing reproducible evolutionary and statistical insights.

Key features of the toolkit include Tajima’s D calculator with Site Frequency Spectrum (SFS), Shannon’s Entropy (H) calculator, Alignment re-formatting tool, HGSV to FASTA conversion, amino acid frequency analysis, FASTA to SEQRES converter, RNA 2D structure alignment tool, and Kurtosis coefficient calculator. While the number of the tools is open to increase, current tools cover an underrepresented spectrum of evolutionary and statistical analyses, enabling researchers to gain additional insights from their genetic sequence data. The toolkit is not only a research tool but also serves as an educational resource. Its interactive interface can help students and new researchers understand the process of alignment analysis and the calculation of relevant evolutionary statistics. By providing clear visualizations and step-by-step explanations, the toolkit bringing theoretical concepts and practical application closer in evolutionary biology.

The Evolutionary Statistics Toolkit is hosted on a user-friendly web interface, designed using HTML for website structure and JavaScript for interactivity and functionality. This allows researchers to navigate through various analytical tools, each performing specific statistical and evolutionary analyses. The toolkit also processes input directly on the client-side, minimizing security risks associated with server-side processing. To ensure transparency and foster community collaboration, the source code for each tool in the toolkit is made available through GitHub links provided on the website. With these links, specific Colab notebooks are also available that increases efficiency for batch analyses. This open-source approach allows users to inspect the code, suggest improvements, or even adapt the tools for their specific research needs. For testing our web toolkit, we specifically utilized genome assemblies from all six hominoid species to create a hominoid-consensus sequence. We conducted statistical tests on hominid canonical aGPCR (adhesion G protein-coupled receptor G1, ADGRG1) transcripts. Through these analyses, we aim to uncover effectiveness of the toolkit for uncovering evolutionary patterns on the biological significance and evolutionary history of these receptors. Furthermore, the toolkit’s capabilities will be demonstrated through various case studies, highlighting its utility in evolutionary genomics research.

In this article, we will describe in detail the functionalities, applications, and validation of each tool within the Evolutionary Statistics Toolkit. We will also discuss the toolkit’s accessibility features, potential improvements, and its overall impact on the field of evolutionary genomics.

## Methods

### Website and Accessibility

The main interface of the Evolutionary Statistics Toolkit is designed for simplicity and functionality. The color gradient uses black and-white coloring for accessibility [4]. The layout is clean and straightforward, with clearly labeled sections and buttons. This approach minimizes the learning curve for new users. The interface employs a clean, hierarchical structure. The main title “Evolutionary statistics toolkit” is prominently displayed at the top, immediately communicating the purpose of the application. The top of the page also features a navigation bar with links to various tools, including SFS & Tajima’s D, Shannon’s H, Alignment Matrix, HGSV to FASTA, Wave-function, Pair Frequencies, and Batch/Source. This allows users to switch between different analyses (Figure 1).

**Figure 1.**
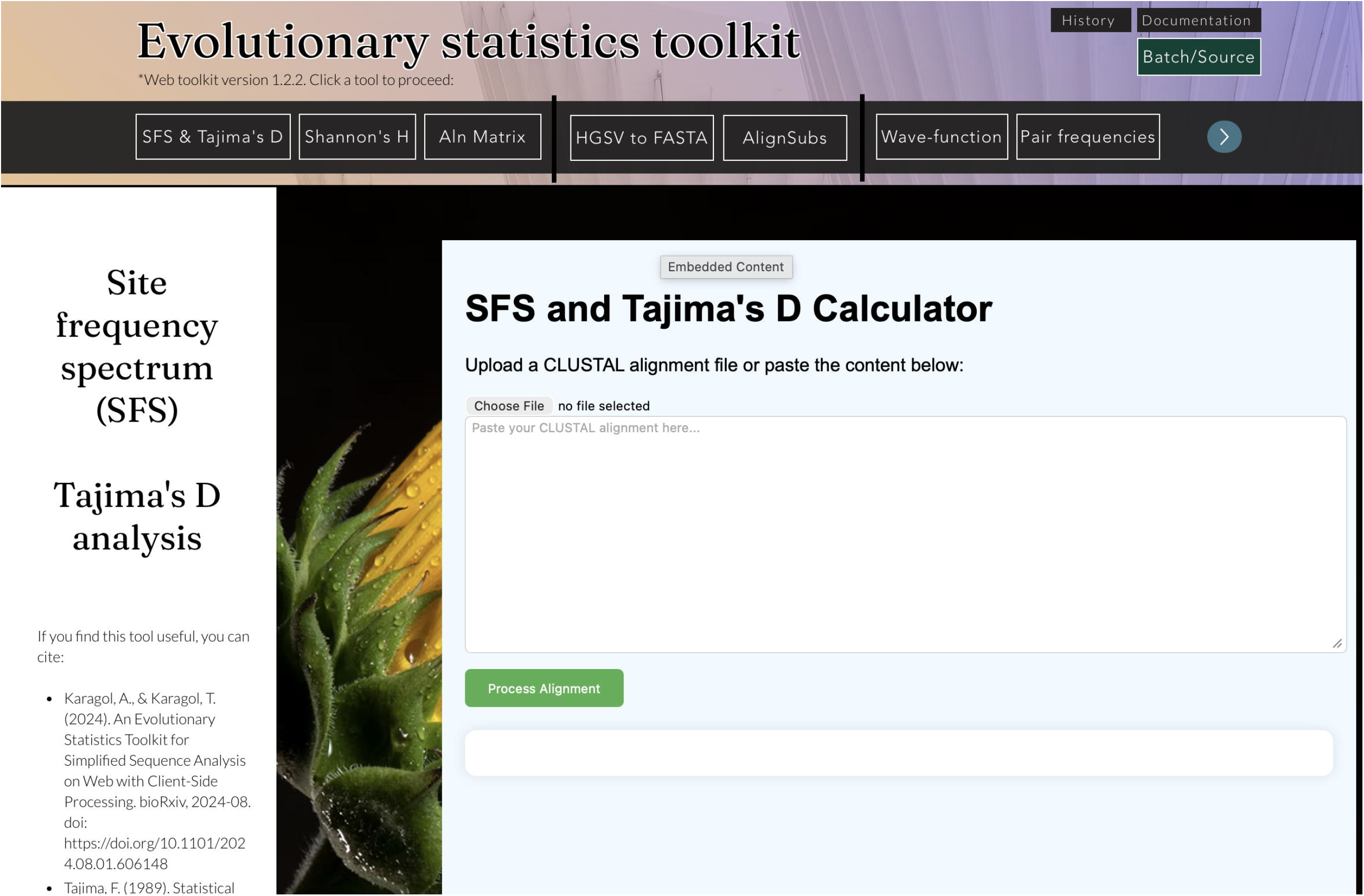
User interface of the website. The top of the page features a navigation bar with links to various tools, including SFS & Tajima’s D, Shannon’s H, Alignment Matrix, HGSV to FASTA, Wave-function, Pair Frequencies, and Batch/Source. Below, the main content area displays the interface for the currently selected tool, in this case, the “SFS and Tajima’s D Calculator”.

The main content area displays the interface for the user selected tool, in this case, the “SFS and Tajima’s D Calculator” (Figure 1). The background images of individual tools are mostly black, with small plants on left side to differentiate each tool. On the left side, the explanation of each tool and proper citations in APA format are located. On the right side, the tools embedded to the website with HTML codes containing each tool algorithm (Table 1). These sections provide a clear input mechanism, allowing users to either upload a CLUSTAL alignment file or paste content directly into a text area. The “Process” buttons are distinctly visible, using color to draw attention. This presentation scheme enables additional tools to be added in future versions. The version page provides the open history of the previous changes on the website, which is accessible in upper left corner. The old versions are also fully functional, with backups of the entire toolkit from that time. This feature helps users and academics with a more robust reproducibility analyses.

**Table 1.**
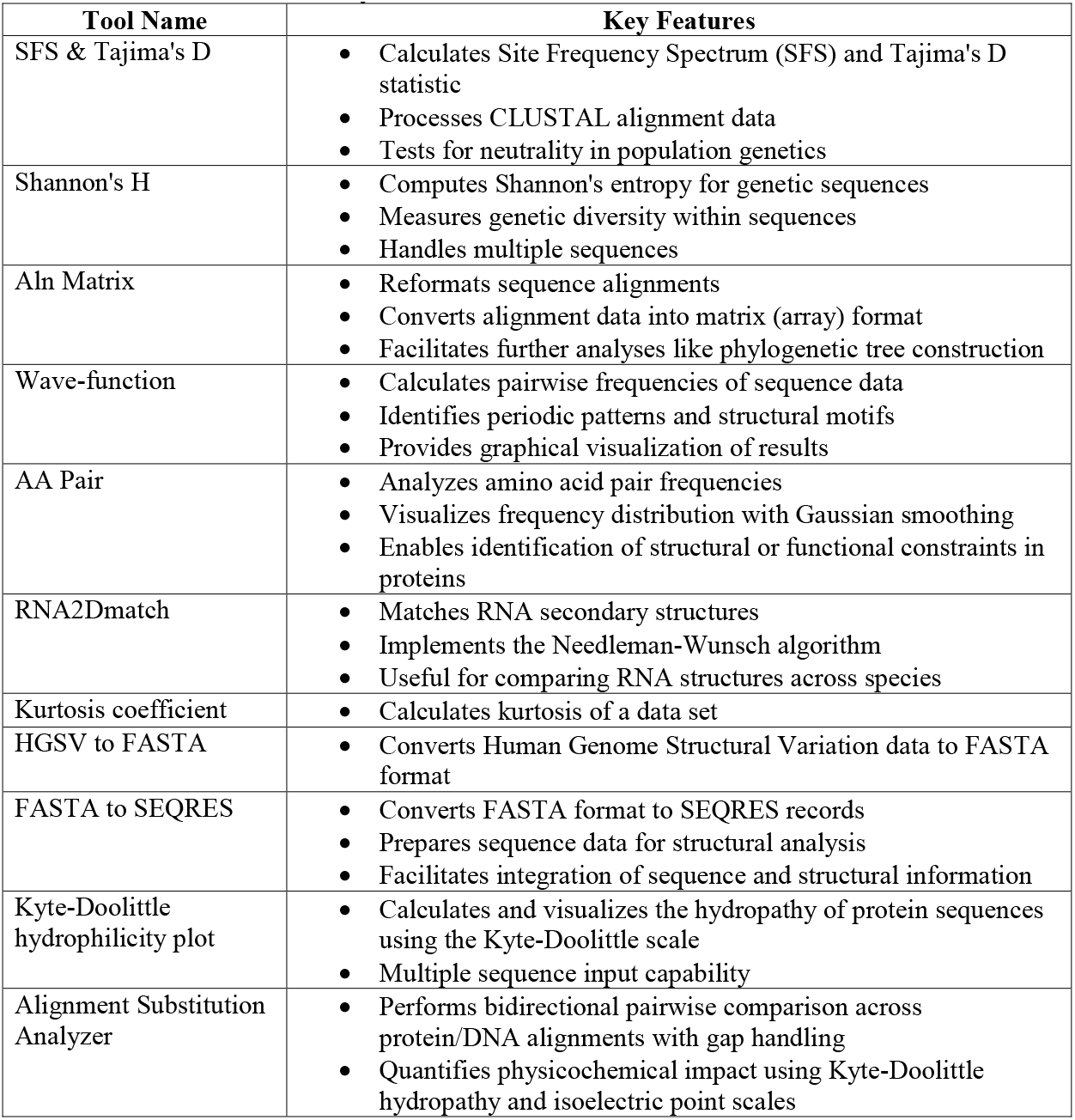
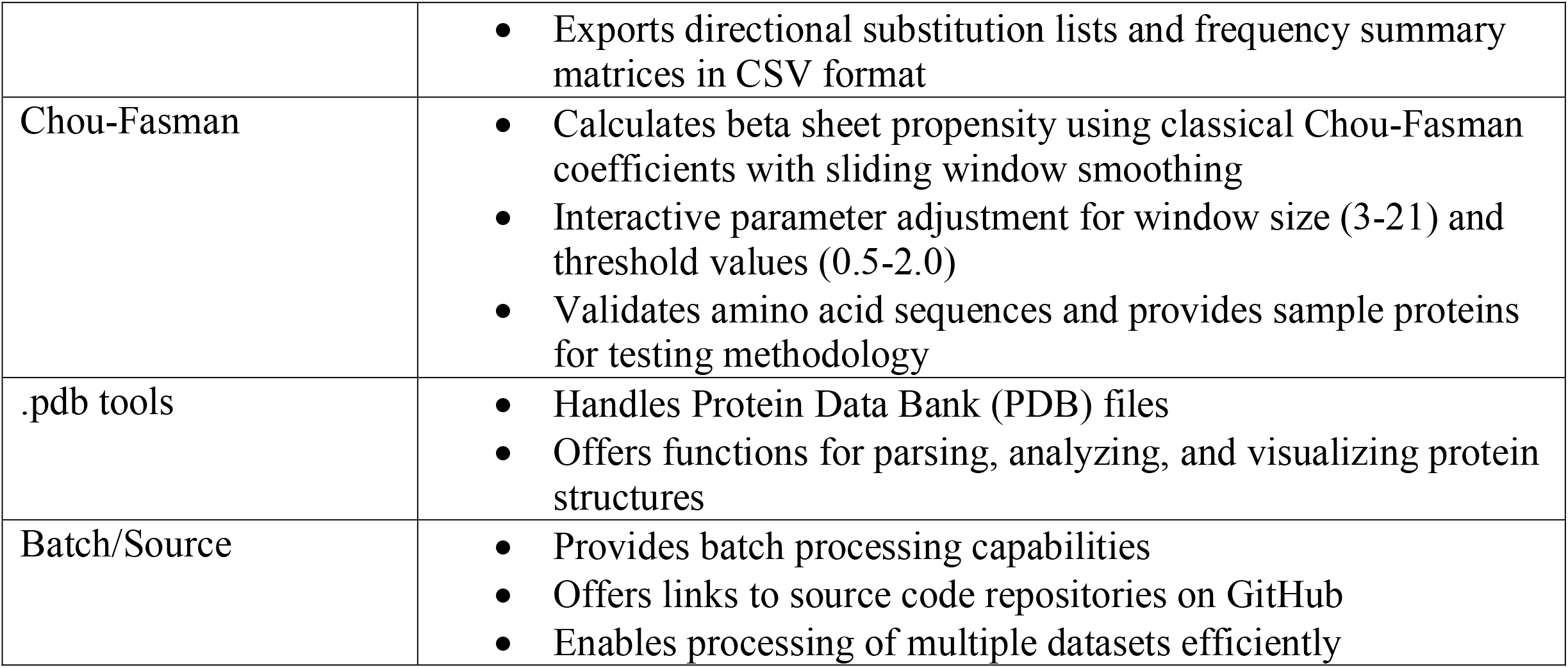
The tools and their key features that are featured on the website.

| **Tool Name** | **Key Features** |
| --- | --- |
| SFS & Tajima's D | • Calculates Site Frequency Spectrum (SFS) and Tajima's D statistic<br>• Processes CLUSTAL alignment data<br>• Tests for neutrality in population genetics |
| Shannon's H | • Computes Shannon's entropy for genetic sequences<br>• Measures genetic diversity within sequences<br>• Handles multiple sequences |
| Aln Matrix | • Reformats sequence alignments<br>• Converts alignment data into matrix (array) format<br>• Facilitates further analyses like phylogenetic tree construction |
| Wave-function | • Calculates pairwise frequencies of sequence data<br>• Identifies periodic patterns and structural motifs<br>• Provides graphical visualization of results |
| AA Pair | • Analyzes amino acid pair frequencies<br>• Visualizes frequency distribution with Gaussian smoothing<br>• Enables identification of structural or functional constraints in proteins |
| RNA2Dmatch | • Matches RNA secondary structures<br>• Implements the Needleman-Wunsch algorithm<br>• Useful for comparing RNA structures across species |
| Kurtosis coefficient | • Calculates kurtosis of a data set |
| HGSV to FASTA | • Converts Human Genome Structural Variation data to FASTA format |
| FASTA to SEQRES | • Converts FASTA format to SEQRES records<br>• Prepares sequence data for structural analysis<br>• Facilitates integration of sequence and structural information |
| Kyte-Doolittle hydrophilicity plot | • Calculates and visualizes the hydropathy of protein sequences using the Kyte-Doolittle scale<br>• Multiple sequence input capability |
| Alignment Substitution Analyzer | • Performs bidirectional pairwise comparison across protein/DNA alignments with gap handling<br>• Quantifies physicochemical impact using Kyte-Doolittle hydropathy and isoelectric point scales |
|  | • Exports directional substitution lists and frequency summary matrices in CSV format |
| Chou-Fasman | • Calculates beta sheet propensity using classical Chou-Fasman coefficients with sliding window smoothing<br>• Interactive parameter adjustment for window size (3-21) and threshold values (0.5-2.0)<br>• Validates amino acid sequences and provides sample proteins for testing methodology |
| .pdb tools | • Handles Protein Data Bank (PDB) files<br>• Offers functions for parsing, analyzing, and visualizing protein structures |
| Batch/Source | • Provides batch processing capabilities<br>• Offers links to source code repositories on GitHub<br>• Enables processing of multiple datasets efficiently |

Links to detailed documentation and support pages are available on the upper left corner, providing users with guidance on using the toolkit effectively. This includes step-by-step instructions, related papers, and contact information for further assistance. In the lower right corner of the website interface, there is a blue chat button. This feature enhances the accessibility and user-friendliness of the toolkit, this support option can increase user confidence in using the toolkit, potentially leading to broader adoption and more effective use of the analytical tools.

### Tools and Functionalities

#### 1. SFS & Tajima’s D

The code provides a simple web interface where users can upload or paste CLUSTAL alignment data. Upon clicking the “Process Alignment” button, the processAlignment() function processes the input data, formats it, and prepares it for further analysis.

Users can either upload a CLUSTAL alignment file or paste the alignment data directly into a text area. The interface includes a file upload button and a large text area for pasting alignment data, making it easy to input genetic sequences (Figure 1). The processAlignment function starts by retrieving the alignment data from the textarea (alignmentInput). It then checks if any data is provided and alerts the user if not. The function splits the input data into lines, removes unnecessary parts (e.g., headers), and counts spaces to determine the sequence structure. It extracts sequence data, calculating the number of sequences and their lengths. This process involves manipulating and reorganizing the alignment data for further analysis. The sequence count info is added to output to check whether the analysis performed correctly. The alignmentMatrix variable holds the final processed alignment data, structured as a 2D array for easy manipulation in the SFS computation. The tool then calculates the Site Frequency Spectrum (SFS) and Tajima’s D statistic [5].

The SFS is a summary of the allele frequencies in a sample of DNA sequences. For n sequences, the SFS is an (n-1)-dimensional vector where the i-th entry represents the number of sites with i derived alleles.

Mathematically, for each polymorphic site j:

SFS[i] = Σ_j I(d_j = i)

where d_j is the number of derived alleles at site j, and I() is the indicator function.

Tajima’s D is a statistical test that compares the number of segregating sites (e.g., singleton, doubletons) to the average number of nucleotide differences [5]. It is used to test for neutrality in a population by determining if the observed DNA sequence variation is consistent with the neutral theory of evolution [5]. This can help detect different selection models.

Tajima’s D = (π - θ_W) / sqrt(V_D)

where:

π is the average number of pairwise differences (estimator of θ based on π)

θ_W is Watterson’s estimator of θ

V_D is the variance of (π - θ_W)

The code calculates these components as follows:

a) π (theta_pi):

π = Σ_i 2i(n-i)SFS[i-1] / (n(n-1))

b) θ_W (theta_w):

θ_W = S / a_1

where S is the number of segregating sites, and a_1 = Σ_i^(n-1) 1/i

c) Variance V_D:

V_D = e_1S + e_2S(S-1)

where e_1 and e_2 are complex functions of n derived in Tajima’s original paper [5]. The implementation uses JavaScript to perform these calculations in the browser.

#### 2. Shannon’s H

In bioinformatics, the ability to quickly and accurately calculate sequence entropy is essential for understanding the variability within sequences, such as DNA, RNA, or proteins. This tool computes Shannon’s entropy, providing a measure of genetic diversity within a sequence [6]. It can be used to quantify the diversity of genetic sequences within a population or the diversity of species within an ecosystem [7].

For a discrete random variable X with possible values {x_1, …, x_n} and probability mass function P(X), the Shannon entropy H(X) is defined as:

H(X) = −∑ P(x_i) * log_2(P(x_i))

where the sum is over all possible values of X.

The calculateSequenceEntropy(sequence) implements this formula. It calculates the probability of each character by dividing its count by the total sequence length. Finally, it computes the entropy using the formula:

entropy = −∑ p_i * log_2(p_i)

where p_i is the probability of each unique character in the sequence.

Users can add more sequences as needed and calculate the entropy of each sequence by clicking the button provided. The calculated entropies are then displayed on the webpage. Performance may degrade if used with very large sequences or a large number of sequences due to the computational complexity of entropy calculation (Please see the Batch/Source section below). Its integration into broader bioinformatics workflows may enhance the analytical capabilities available to researchers, during the study of genetic diversity and evolutionary biology.

#### 3. Aln Matrix

This tool is a utility for reformatting sequence alignments for compatibility with pairwise analysis tools and simplifies codes for pairwise analysis by making arrays of sequences. The code allows users to interact with the webpage by uploading or pasting sequence alignment data, processing this data, and then downloading the results.

The JavaScript code is embedded within <script> tags and contains the processAlignment() function that handles the sequence alignment data processing. It essentially transforms a vertically aligned set of sequences into a horizontally aligned set, while preserving the alignment relationships. This is achieved through the careful calculation of indices and the creation of a 2D matrix.

The index calculation (i * a + i) + c is particularly interesting. It is designed to jump between sequences in the original format to create a continuous string of aligned sequences. This formula ensures that characters from each sequence are picked in the correct order to maintain the alignment. By converting alignment data into a matrix (array) format, it simplifies further analyses such as phylogenetic tree construction, sequence motif identification, and comparative genomics. It is used in multiple sequence alignments to identify regions of similarity that may indicate evolutionary relationships among the sequences.

#### 4. Wave-function

The “wave function” in this context is not a quantum mechanical wave function, but rather a plot of entropy values:

WaveFunction = {(i, H(i)) | i = 1 to L}

where L is the length of the sequences.

Shannon entropy, calculated for each position i:

H(i) = −∑[f(a,i) * log_2(f(a,i))]

where H(i) is the entropy at position i, and the sum is taken over all characters a present at that position [6].

The entropy values are plotted against sequence position, creating a graph that visualizes the variability along the sequence:

x-axis: Position i (1 to L)

y-axis: Entropy H(i).

The code uses Chart.js for visualizing the sequence data. The results section includes elements for displaying calculated data and visualizations. This functionality is beneficial for bioinformatics toolkits as it aids in the visualization and interpretation of sequence data, producing insights into evolutionary relationships.

#### 5. AA Pair frequencies with Gaussian Smoothing

By providing a visual representation of these frequencies, this tool enables researchers to identify patterns, biases, or conservation in amino acid pairings, which can be indicative of structural or functional evolutionary constraints. The representation with Gaussian smoothing offers a continuous view of the frequency distribution, potentially revealing trends that might be less apparent in discrete frequency plots.

The core mathematical concept used here is Gaussian smoothing, which is applied to the raw frequency data. The smoothed wave function is defined as:

Ψ(x) = Σ A_i * exp(-((x – x_i)^2^ / (2σ^2^)))

where:

Ψ(x) is the wave function value at position x

A_i is the amplitude (frequency) of the i-th amino acid pair

X_i is the position of the i-th amino acid pair

σ is the standard deviation of the Gaussian function (set to 1 in this code)

For each point x in the dataset, the wave function value is calculated by summing the contributions from all other points, weighted by a Gaussian function:

calculateWaveFunction(x, data) = Σ_i data[i].y * exp(-((xIndex - i)^2^ / (2σ^2^)))

where xIndex is the index of x in the dataset, and data[i].y is the frequency of the i-thpair.

The plotWaveFunction() utilizes Chart.js to create a line chart representing the wave function. downloadResults() allows users to save the processed data in JSON format. The JSON output format allows for easy integration with other bioinformatics tools and scripts for further analysis.

#### 6. RNA2Dmatch

This tool matches RNA secondary structures of 2 input RNAs, which is useful in evolutionary studies to analyze structural conservation across species. This functionality is added to toolkit for comparing RNA structures, identifying conserved structural elements, and understanding evolutionary relationships between RNA molecules. While there is a room for optimization and feature development, this tool represents a solid foundation for web-based RNA structure comparison. The alignStructures() function retrieves user input, validates it, and calls the alignment algorithm. The needlemanWunsch() implements the Needleman-Wunsch algorithm for global sequence alignment [8]. By doing this the codes incorporates a scoring system for matches, mismatches, gaps, and a traceback procedure to reconstruct the optimal alignment. Then, displayAlignment() formats and displays the alignment results, including: sequence position numbering and line-by-line output for readability.

This tool can be integrated into various bioinformatics workflows as a quick comparison tool for RNA structures predicted by different algorithms. The Needleman-Wunsch algorithm has a time complexity of O(mn), where m and n are the lengths of the input sequences [8]. For very long RNA structures, this could lead to performance issues in the browser. This tool would be used to compare RNA secondary structures to identify common motifs or structural similarities, which can be crucial for understanding RNA function, predicting RNA behavior, and studying evolutionary conservation of RNA structures.

#### 7. Kurtosis coefficient

Statistical measures are helpful for understanding the distribution and patterns of genetic variation. One such measure is kurtosis, which describes the shape of a probability distribution, particularly its “tailedness” relative to a normal distribution [9]. This tool calculates the kurtosis coefficient of a data set. It includes functionality for data input (either via file upload or manual entry), kurtosis calculation, data visualization through a line chart, and the ability to download results in both text and PDF formats. The application utilizes JavaScript for calculations and interactivity and integrates Chart.js for graphing and jsPDF for PDF generation. In evolutionary biology, the kurtosis coefficient can be used to analyze the distribution of sequence variants. High kurtosis indicates more outliers, which can signify rare evolutionary events or selection pressures.

#### 8. HGSV to FASTA

This converter tool transforms Human Genome Structural Variation (HGSV) data into FASTA format, based on wild-type protein/nucleotide sequences and variant notations in HGVS format [10,11]. The conversion to FASTA format makes the structural variant data compatible with many bioinformatics tools for further analysis, such as sequence alignment, variant calling, and genetic analysis. This tool is particularly useful for researchers and bioinformaticians working with protein sequences to study the effects of missense variants (Figure 2). It can also be easily installed on a local computer using pip, provided Python 3.x is available, with the following command: pip install HGVStoFASTA.

**Figure 2.**
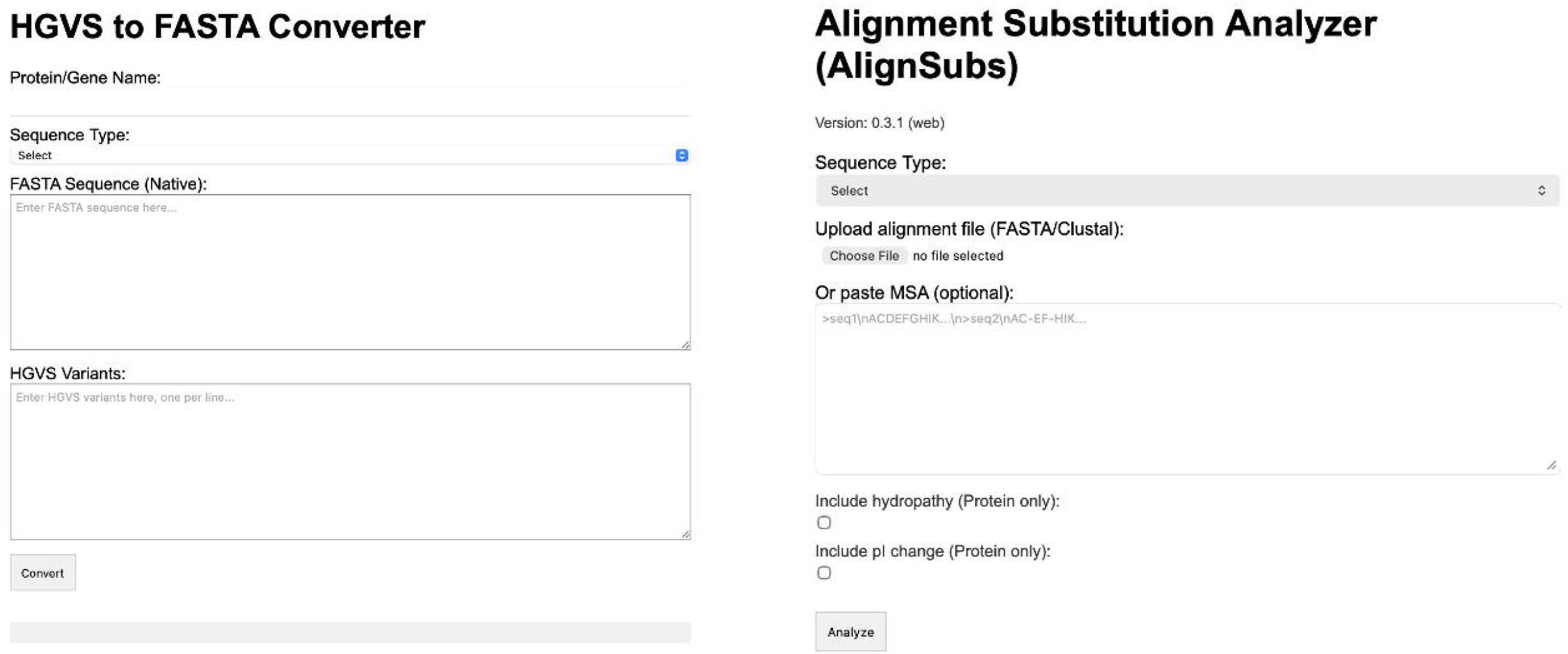
HGSV to FASTA converter and AlignSubs. This converter tool transforms Human Genome Structural Variation (HGSV) data into FASTA format, based on wild-type protein/nucleotide sequences and variant notations in HGVS format. AlignSubs tool performs bidirectional pairwise comparison across protein/DNA alignments with gap handling.

#### 9. FASTA to SEQRES

This tool is used to prepare sequence data for structural analysis in protein databases, for the integration of sequence information with protein modeling and analysis. The FASTA format is widely used for representing nucleotide or peptide sequences, while SEQRES records are essential components of Protein Data Bank (PDB) files, providing a standardized representation of macromolecular structures [12].

#### 10. Kyte-Doolittle hydrophilicity plot

The tool for generating hydropathy score plots measures the hydrophobic or hydrophilic nature of amino acid residues along the protein sequence [13,14]. The Kyte-Doolittle hydropathy scale assigns a hydropathy value to each amino acid based on its physicochemical properties [13]. For each residue in the sequence, the generated plot shows the hydropathy value. Positive values indicate hydrophobic regions, while negative values indicate hydrophilic regions. Peaks above a certain threshold for a stretch of residues may indicate potential transmembrane regions. Furthermore, our script offers batch processing capabilities, saving time and effort when analyzing multiple sequences, which is scalable to handle large datasets, limited only by local hardware. This tool can also be installed via pip, utilizing the following command: pip install hydrophilicity-plot. It supports both manual sequence input and file-based sequence input.

#### 11. AlignSubs (Alignment Substitution Analyzer)

This tool provides directional substitution lists and frequency summary matrices for an alignment (Figure 2). Input sequences undergo character-level validation against defined alphabets: [ACGTU-] for nucleic acids where gaps are represented by hyphens. The algorithm implements bidirectional pairwise comparison across sequences, generating comparison matrices for comprehensive substitution pattern analysis. Sequence type validation ensures data integrity through user-confirmed character set compliance, with optional continuation for ambiguous input containing non-standard residues. For protein sequences, substitution events are characterized using two established scales: the Kyte-Doolittle [13] hydropathy index [−4.5, 4.5] and isoelectric point values pI [2.77, 10.76] for each amino acid. Property changes are calculated as Δ parameters: ΔH = H(R_j) - H(R_i) and ΔpI = pI(R_j) - pI(R_i) for each substitution R_i → R_j. Gap positions are excluded from substitution calculations. The hydropathy scale ranges from highly hydrophobic (Ile: 4.5) to highly hydrophilic (Arg: −4.5), while pI values span from acidic (Asp: 2.77) to basic (Arg: 10.76) residues. The framework generates two complementary CSV outputs: directional substitution lists containing position-specific changes with counts and property deltas, and summary matrices aggregating substitution frequencies across all sequence pairs. This tool can also be installed via pip, utilizing the following command: pip install AlignSubs.

#### 12. Chou-Fasman SIMPLE

The Chou-Fasman method employs empirically derived amino acid propensity coefficients P(R_i) for β-sheet formation, where each residue type exhibits a normalized frequency of occurrence in β-sheet conformations [15]. The algorithm implements a sliding window approach across protein sequences of length n, calculating local β-sheet density β(i) at position i using a convolution.The computed propensity is assigned to the window centroid at position i + ⌊w/2⌋ + 1, ensuring symmetric neighborhood representation around each analyzed position.

The propensity matrix includes values ranging from P(Glu) = 0.37 (strong β-sheet breaker) to P(Val) = 1.70 (β-sheet former), with the theoretical baseline P = 1.0 representing statistical equivalence to random occurrence [15]. Regions satisfying β(i) > τ, where τ is the user-defined threshold are classified as potential β-sheet forming segments. The algorithm maintains computational complexity O(n·w) with linear memory scaling, processing sequences through validation and enforcing minimum sequence length constraints of 10 residues to ensure statistical significance of windowed analyses. The computational implementation generates continuous propensity profiles β(i) mapped against residue positions, with threshold visualization rendered as a horizontal reference line. Multi-sequence comparative analysis is supported through overlay plotting. The SIMPLE version of the tool allows manuel entering of multiple protein sequences, without any spesific alignment formats (Figure 3).

**Figure 3.**
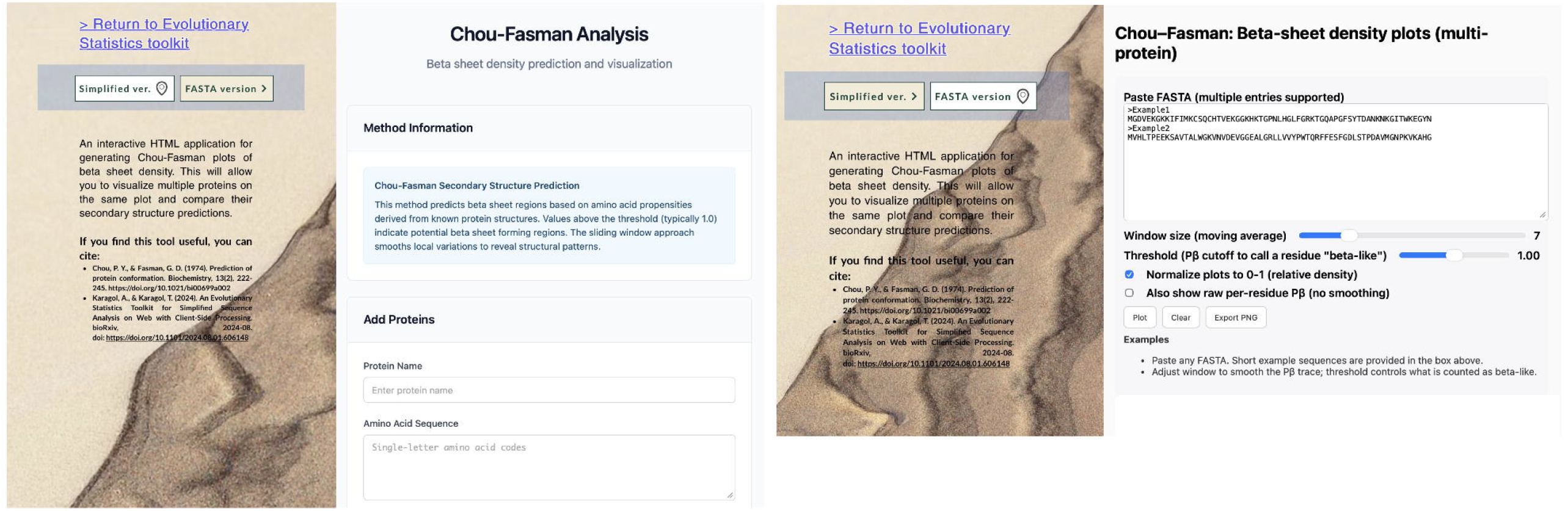
User interface of the Chou-Fasman tool. This tool calculates beta sheet propensity using classical Chou-Fasman coefficients with sliding window smoothing and provides interactive parameter adjustment for window size (3-21) and threshold values (0.5-2.0).

#### 13. Chou-Fasman FASTA

In this version, multiple protein sequences are processed simultaneously through FASTA parsing with automatic header extraction and sequence validation using the amino acid alphabet. The algorithm supports optional min-max normalization: β_norm(i) = (β(i) - β_min)/(β_max - β_min), transforming propensity profiles to [0,1] intervals for comparative analysis across proteins with different baseline propensities. Binary classification of β-sheet regions employs a threshold function: S(i) = 1 if β(i) ≥ τ, 0 otherwise, where τ represents the user-defined cutoff (range: 0-2, default: 1.0). Quantitative β-sheet density metrics are computed as: D_β = |{i: β(i) ≥ τ}|/n, representing the fraction of residues exceeding the threshold criterion.

The algorithm implements a sliding window convolution across protein sequences of length n, calculating local β-sheet density β(i) at residue position i using: β(i) = (1/w) × Σ_{j=i-⌊w/2⌋}^{i+⌊w/2⌋} P_β(R_j), where w represents the window size (adjustable: 1-31 residues, default: 7) and P_β(R_j) denotes the propensity value for residue (R_j). The moving average implementation provides symmetric neighborhood representation around each position, with boundary effects handled through adaptive window sizing near sequence termini. Multi-sequence comparative analysis is supported through overlay plotting with distinct chromatic encoding for each protein sequence.

#### 14. pdb tools

These tools handle PDB (Protein Data Bank) files, which are implemented for combing evolutionary analyses with structural data. These tools could be used for structural bioinformatics, enabling researchers to study protein conformations, analyze molecular interactions, and increase visualization of structural models (Figure 4). The B-factor (or temperature factor) tool adds B-factors in PDB files. B-factors indicate the specific coloring codes for structures. The Chain Swapper tool facilitates the exchange of polypeptide chain naming within a PDB structure, making it particularly useful for representing data on a single chain in heterodimers. An extended version of the Chain Swapper allows for the exchange of three chains simultaneously, enabling more complex structural visualizations. Another tool combines RNA structures from separate PDB files into a single PDB file, which is valuable for tools that require a combined PDB file for RNA and protein structures.

**Figure 4.**
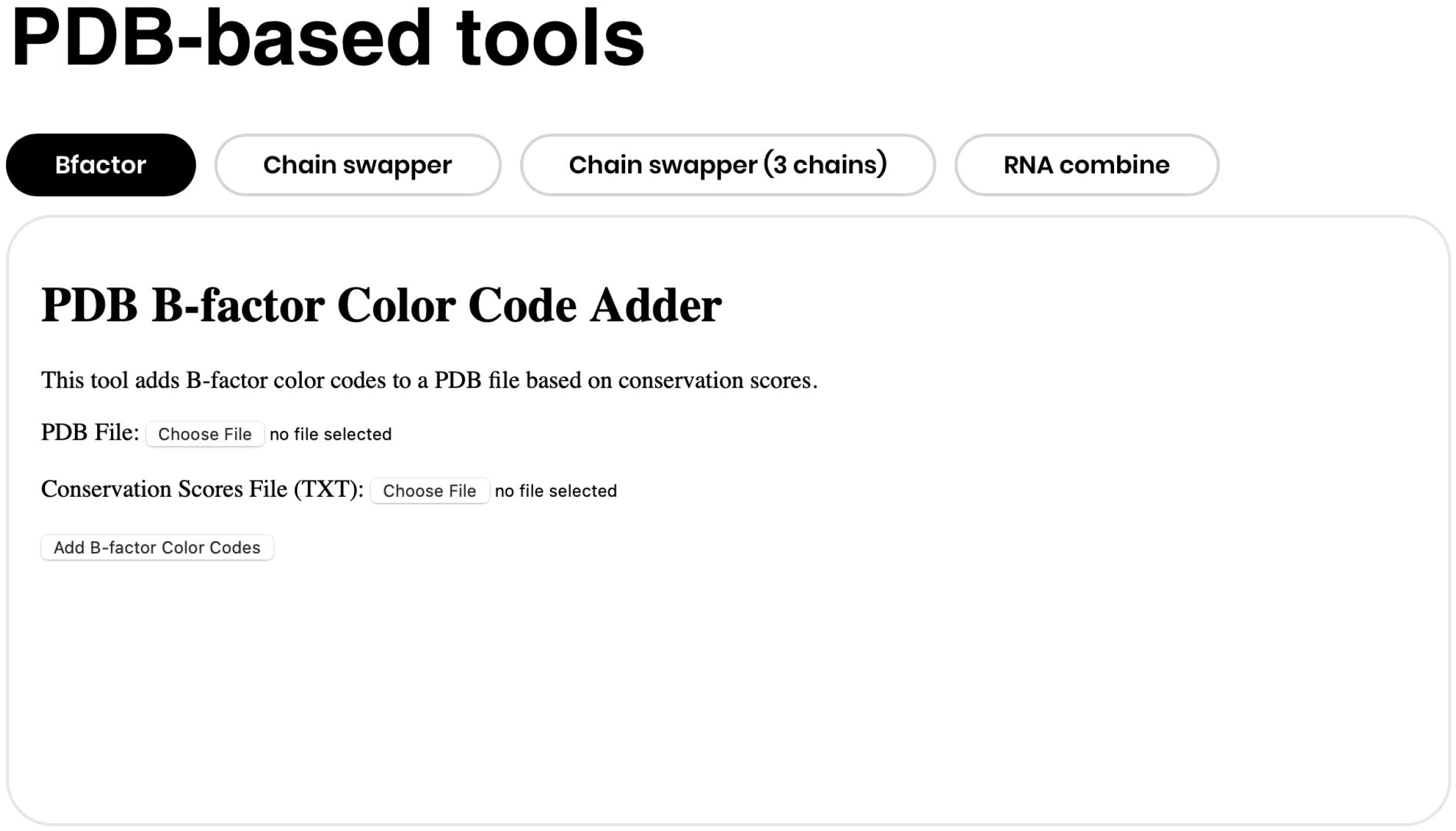
Tools for.pdb files. The B-factor (or temperature factor) tool analyzes and modifies B-factors in PDB files. B-factors indicate the specific coloring codes for structures. Chain swapper tool facilitates the exchange of polypeptide chains between different PDB structures. RNA combine tool combines a protein PDB file and an RNA.

## Results

### Validation

We validated the toolkit using our previous research data. Some tools like SEQRES format converter previously demonstrated its efficacy in uncovering evolutionary patterns and genetic diversity [16]. The SFS & Tajima’s D tool successfully identified selection signatures. The alignment re-formatting and HGSV to FASTA tools could be streamline the data preparation workflows, while the pair analysis tools may reveal significant periodic patterns in genomic sequences. The CSS styles ensure that the page has a consistent and appealing look. The body has a light blue background, text areas are full-width, and buttons have a distinct color for visibility. The entire website was also checked with color blindness simulator tool on the Wix, to provide reliable accessibility. The evolutionary statistics toolkit hosted on Wix, that offers security features including SSL encryption to protect data in transit, DDoS protection to safeguard against denial-of-service attacks, and regular security updates to address emerging threats.

The toolkit processes alignment data on the client side, which may be slow for large datasets. Accordingly, the Batch/Source segment offers functionalities for processing multiple data sets (batch processing) and provides links to the source code repositories (collar links) on GitHub. This allows users to run the tools on multiple data sets more efficiently on Colab or Python, which is essential for large-scale studies and analyses [17]. Batch processing can save time and ensure consistency. By providing access to the source code, users can understand how the tools work, verify the methods used, and potentially customize the tools to better fit their specific needs

The toolkit could be integrated with existing bioinformatics tools and libraries (e.g., Biopython) to leverage their additional functionalities and provide a more comprehensive analysis. Another future application is the optimization of the processing algorithm to handle larger datasets more efficiently. Improving the UI would also be beneficial. Future updates may include progress indicators, detailed instructions, and more user-friendly input methods (e.g., drag-and-drop for file uploads).

### Comparison with other tools

There is no available user-friendly toolkit for SFS and Tajima’s D calculation. On the other hand, we compared the calculation of the toolkit with existing codes for Tajima’s D. For the input, we first generated an Clustal alignment file containing hominid canonical aGPCRs (adhesion G protein-coupled receptor G1, ADGRG1), their cDNAs were obtained from Ensemble genomes [18]. Several human diseases have been linked to variants of aGPCRs, a single mutation in the human ADGRG1 (GPR56) gene has been identified as a cause of brain cortical malformation [19,20]. Additionally, this protein holds evolutionary significance, as all nine families of aGPCRs found in humans were already established at the onset of vertebrate evolution [21]. For this test of our web toolkit, we specifically chose genome assemblies from all six hominoid species (GRCh38, gorGor4, Pan_tro_3.0, panpan1.1, Susie_PABv2, and Nleu_3.0) for a hominoid-consensus sequence. Namely: the transcripts of ENST00000562631.7, ENSGGOT00000008484.3, ENSPTRT00000062018.4, ENSPPAT00000032022.1, ENSPPYT00000008695.3, ENSNLET00000004000.2. Clustal Omega was utilized for sequence alignment generation [22,23].

Site Frequency Spectrum (SFS): 395, 125, 296, 2066, 1350 (Supplementary Figure S1). This variational data results in a Tajima’s D value of 0.2616. While further analysis is a necessity, the slightly positive value (0.2616) is close to zero, which suggests that the sequences analyzed are likely evolving neutrally, if confirmed by complementary analyses [5]. There might be a slight indication of balancing selection or population subdivision, but it is not a strong signal. It is important to note that, these values are more meaningful in comparison with other aligned sequences [5]. Nevertheless, this tool provides insights into the evolutionary history of these hominoid species by comparing specific genetic sequences across them.

We compared this result with using user input option; the toolkit produces the same result as uploading the alignment file. For further validation and improvements, the R codes for the SFS are also available in the GitHub links provided on the website. While we cannot identify a similar web interface, another source code for Tajima’s D were also returns same results from the SFS data we calculated via our web toolkit (https://gist.github.com/mt1022/32e54792dbf4df40da2f2a4b87d218c3). The fragmented flow result in difficulties since the SFS calculation was also unavailable and the SFS data must be independently obtained for this code. The existing codes for a similar calculation required excessive coding skills and no GUI to interface (https://github.com/WhalleyT/tajima).

For Shannon’s entropy calculation, we utilized the consensus generated from percentage identities along with previously utilized transcripts (Supplementary Figure S2). In this analysis, the entropy values did not show major disparities, indicating that the diversity of the sequences across the hominoid species was relatively uniform (Supplementary Table S1). For the Percentage Identity Hominid Consensus, the value was H= 2.0526 (Supplementary Table S1). The entropy measures the randomness or uncertainty in the sequence [6].

For the array format, the website was tested with the Clustal Omega alignment that was also utilized for sequence alignment generation. The code automatically generated and downloaded the processed array format containing sequences for the six hominid transcripts. Our array format used on HTML tools on VS Code and R GUI with a simple copy-paste. The Github link provides Colab notebook for batch analysis which also results in the same output. This array format also used in another tool on the website: The Gaussian smoothing equation is used to smooth the data, making it easier to identify trends and patterns. In the output graph, the x-axis labels represent various pairs of amino acids (e.g., AA, AC, AG, AT, etc.). These pairs are crucial for understanding the sequence composition and potential functional motifs within the ADGRG1 sequences. The y-axis values show the frequency or count of each amino acid pair in the input sequences. Higher values suggest that certain pairs are more prevalent in the sequences analyzed. The peaks in the plot indicate amino acid pairs that occur more frequently. For example, the pair “CC” has a high value around 3000, indicating it is very common in the sequences (Supplementary Figure S3). Understanding which pairs are more common can help in designing experiments to mutate specific regions and study their effects on protein function.

The RNA sequences were canonical Human and Gorilla ADGRG1 (GRCh38, gorGor4) cDNAs that were retained from Ensemble genomes, which were also utilized for Tajima’s D calculation. we compared this tool with the RNAFold package 2.0 predicted RNA secondary structures as inputs for our web toolkit [24,25]. Based on the 2D structure alignment provided, a high degree of structural similarity between the human (Seq1) and gorilla (Seq2) ADGRG1 RNA sequences were identified (Supplementary Figure S4). There are numerous regions where the structure is highly conserved between the two species. The region from position 421 to 540 shows almost perfect structural conservation. Another well-conserved region can be seen from position 901 to about 1080. The 3’ end of the sequence (approximately positions 4261 to 4537) also shows high structural similarity. There are also differences in the middle section, particularly around positions 2700-3000. Interestingly, 3’end of this region also showed a higher entropy (Supplementary Figure S3). While the structural alignment does not directly provide information about Tajima’s D values, it complements the analysis by offering insights into structural conservation. Areas with high structural conservation might be under purifying selection, which could be reflected in Tajima’s D calculations.

The sequence conversion utilities were compared with two independent websites: Bugaco (sequenceconversion.bugaco.com) and the ZhangGroup (zhanggroup.org/pdb2fasta/). While both tools have a substantial amount of conversion capabilities, these websites could not handle FASTA to SEQRES conversion. Another website we tried (www.bioinformatics.org/sms2/one_to_three) could convert a FASTA format to three-letter notations but without spaces between characters (Supplementary Figure S6). This website also lacks upload and downland features. The sequence of ADGRG1 accessed utilizing Uniprot (ID: Q9Y653) [26]. Our tool converted the FASTA sequence to SEQRES format (Supplementary Figure S5). Our previous VBA based SEQRES format converter demonstrated its efficacy in uncovering evolutionary patterns and genetic diversity [15]. The HGSV to FASTA converter also successfully implemented the HGSV formatted variation info to sequences in FASTA format. FASTA formats for p.Met1Ala and p.Thr2Ala variants of ADGRG1 were provided (Supplementary Figure S7, Figure S8).

Our kurtosis calculator handles more than 500 data values compared to other web services (Supplementary Figure S9). (www.socscistatistics.com/tests/skewness/). Another service could calculate from more values but without graphical output (Supplementary Figure S9). (exploringfinance.com/kurtosis-calculator/). Many other services are also lacking a proper visualization and downloading feature (Supplementary Figure S10). The kurtosis coefficient calculator tested with segments of Pi digits ranging from 500 to 10,000, with each segment containing 10 digits. The kurtosis of this dataset was calculated −1.1501 by both tools (Supplementary Figure S11).

We compared the hydrophilicity plot tool (Kyte-Doolittle Analysis) with the the ExPASy ProtScale that runs on a web server [27]. While the ExPASy ProtScale tool is a valuable and user-friendly web-based resource, our Kyte-Doolittle script offers several significant advantages, particularly for researchers working with multiple sequences or requiring more customized analyses. The ability to process multiple sequences and creating batch plots in a single pdf file makes our script a powerful tool for in-depth protein sequence analysis (Supplementary Figure S12). For the analysis, the FASTA file containing the sequence of ADGRG1 was accessed utilizing Uniprot (ID: Q9Y653) [25]. The generated plot showed the hydropathy scores for each individual residue with positive values indicate hydrophobic regions, while negative values indicate hydrophilic regions. The negative dips suggest certain regions (e.g., around residues 160, 300, and 500) more exposed to water or hydrophilic areas, possibly involved in ligand binding or extracellular functions (Supplementary Figure S13). Additionally, the local execution and open-source nature of our script provide greater flexibility and potential for customization to meet specific research needs.

## Conclusion

The “Evolutionary Statistics Toolkit” offers an integrated solution for genetic sequence analysis, addressing the gaps in traditional workflows. By combining multiple analytical tools into a single platform, it enhances accessibility and efficiency for researchers. Future developments will focus on expanding tool functionalities and incorporating additional statistical measures to further enhance the toolkit’s applicability.

The “Evolutionary Statistics Toolkit” provides researchers with an intuitive platform for comprehensive sequence analysis. Its integration of multiple analytical tools facilitates a deeper understanding of genetic data, producing new insights into evolutionary dynamics.

## Availability

The toolkit is freely available for academic and non-commercial use. The data generated for the comparisons and validation is available at https://github.com/karagol-alper/evolutionary-toolkit. Source codes of each tool made available in Github with the open GPL-3 license. The GNU General Public License (GPL) is a free, copyleft license for software and other kinds of works. The toolkit is freely accessible on: https://www.alperkaragol.com/toolkit

## Acknowledgments

We thank the research community for valuable feedback during the testing.

