## Supplementary Figure for "An Evolutionary Statistics Toolkit for Simplified Sequence Analysis on Web with Client-Side Processing"

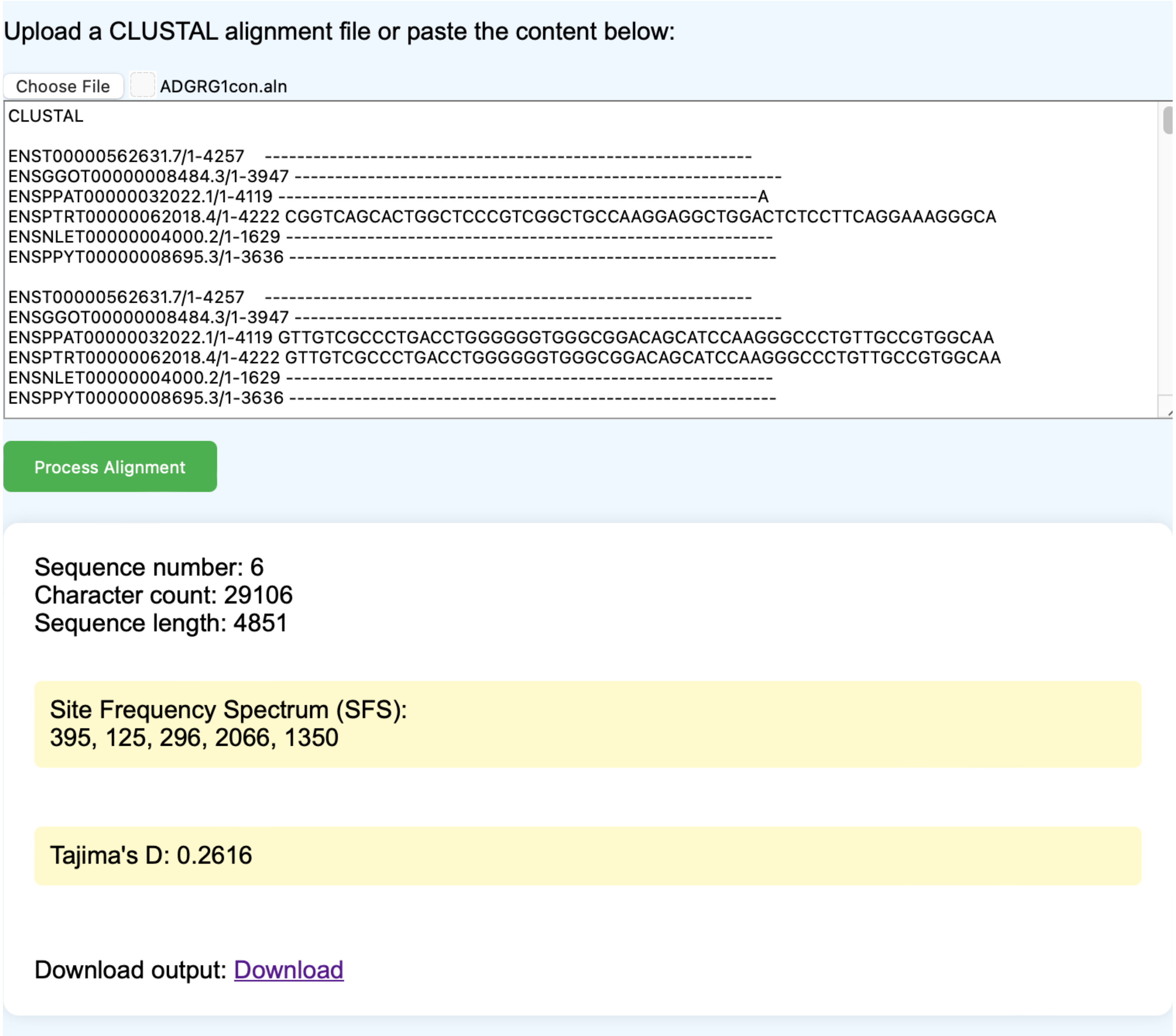


**Figure S1. SFS & Tajima's D calculation.** The values calculated for hominid aGPCRs (adhesion G protein-coupled receptor G1, ADGRG1) cDNAs. We specifically selected genome assemblies of all six hominoid species (GRCh38, gorGor4, Pan_tro_3.0, panpan1.1, Susie_PABv2, and Nleu_3.0) for a hominoid-consensus sequence. Namely: the transcripts of ENST00000562631.7, ENSGGOT00000008484.3, ENSPTRT00000062018.4, ENSPPAT00000032022.1, ENSPPYT00000008695.3, ENSNLET00000004000.2

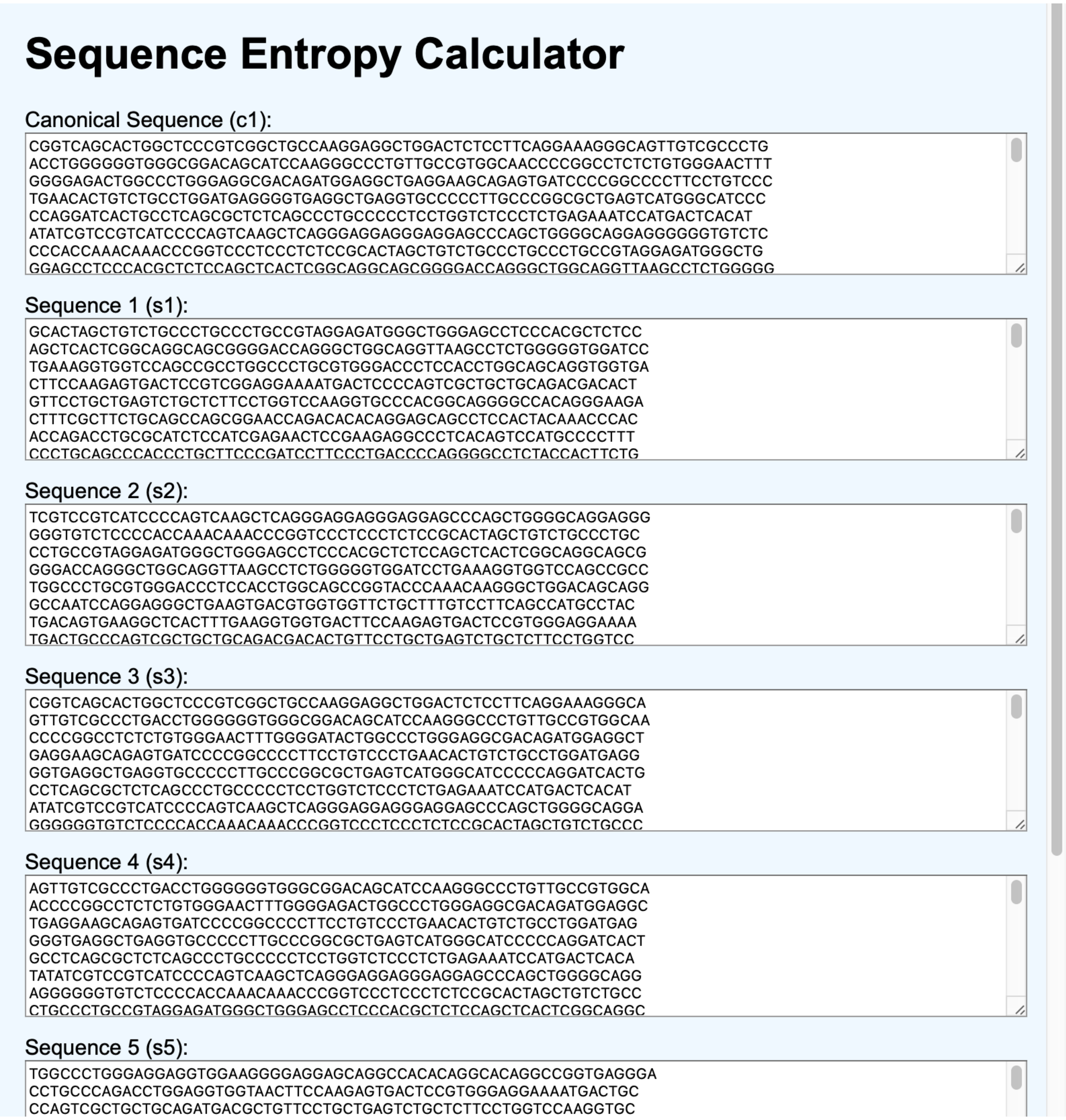


**Figure S2. Input interface of the Shannon’s H calculation tool for ADGRG1 cDNAs.**

**Table S1. Shannon's H values for sampled transcripts, calculated by the toolkit.**

| Species | Transcript ID | Shannon's Entropy |
|---|---|---|
| Hominid Consensus | Clustal aligned consensus | Canonical (c1): 2.0526 |
| Homo sapiens (human) | ENST00000562631.7 | s1: 2.0657 |
| Gorilla gorilla gorilla (western lowland gorilla) | ENSGGOT00000008484.3 | s2: 2.0595 |
| Pan troglodytes (chimpanzee) | ENSPTRT00000062018.4 | s3: 2.0548 |
| Pan paniscus (bonobo) | ENSPPAT00000032022.1 | s4: 2.0560 |
| Pongo abelii (Sumatran orangutan) | ENSPPYT00000008695.3 | s5: 2.0593 |
| Nomascus leucogenys (northern white-cheeked gibbon) | ENSNLET00000004000.2 | s6: 2.0523 |

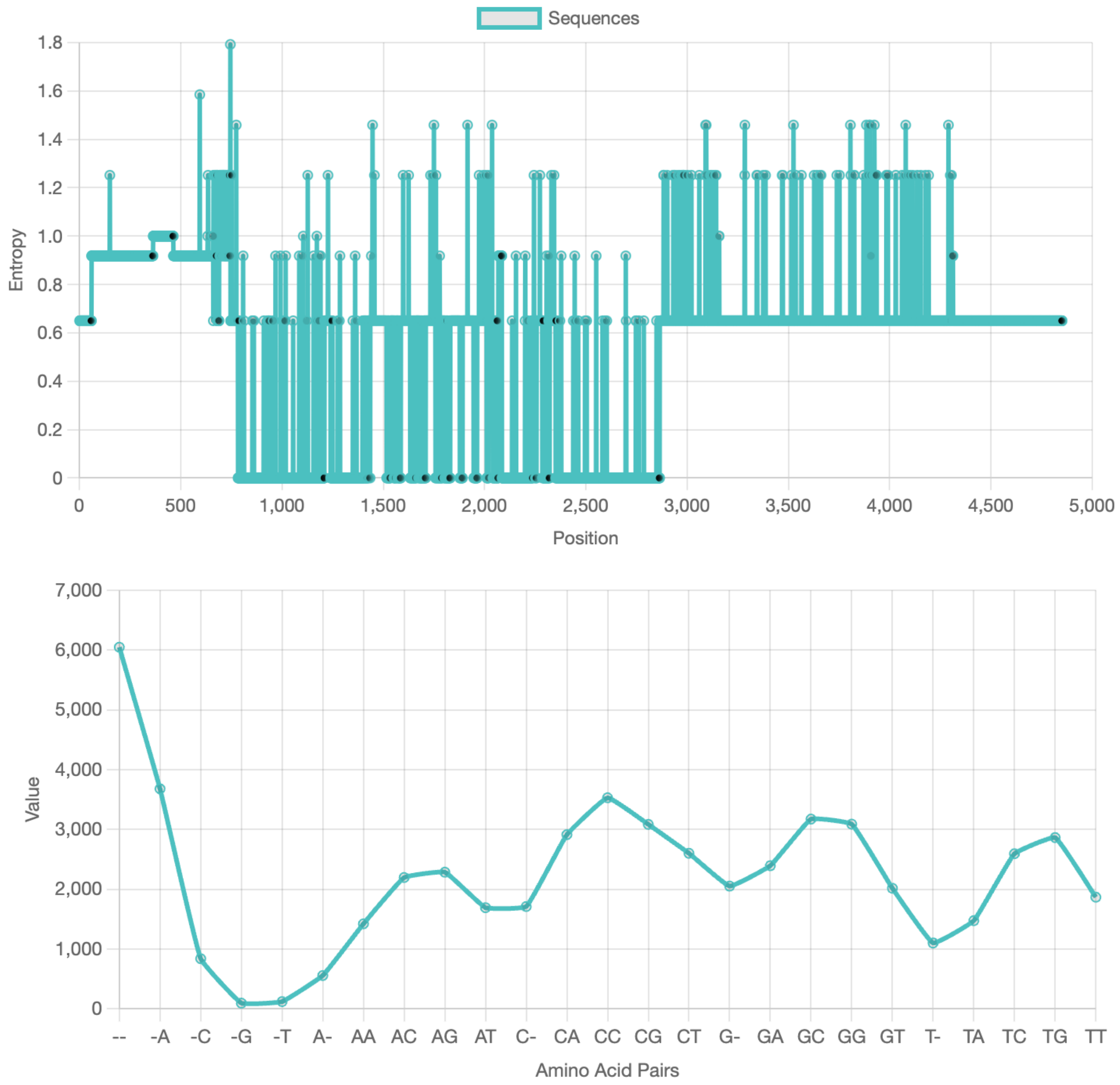


*Gaussian Smoothing Equation:* $\Psi(x) = \Sigma A\_i * exp(-((x - x\_i)\^2 / (2 * \sigma\_i\^2)))$

**Figure S3. Plots from the pairwise analysis output for hominid ADGRG1 cDNAs.**

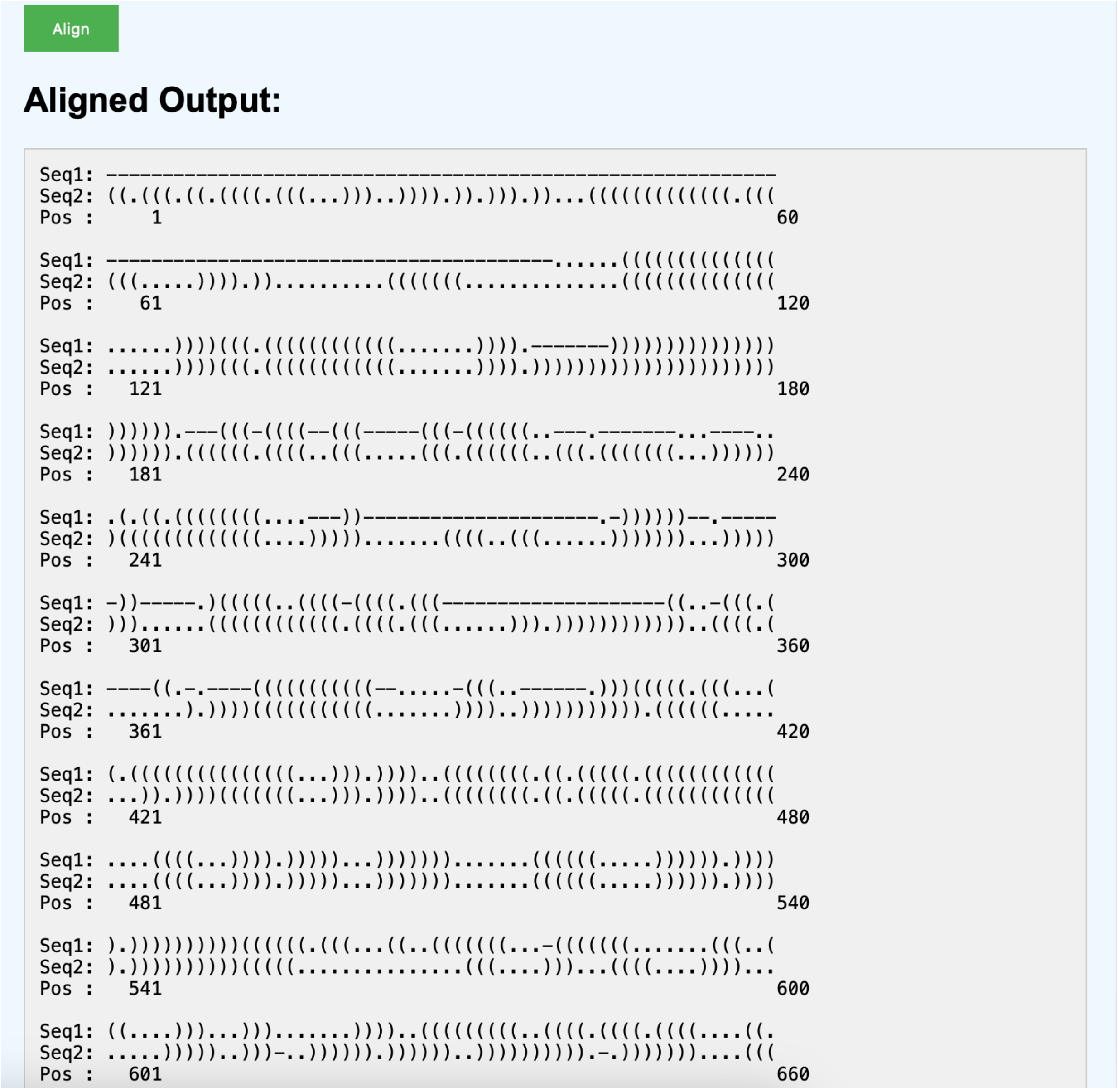


**Figure S4. Output of the RNA2DMatch tool for hominid ADGRG1 cDNAs.** The 2D structures obtained from RNAFold application. The transcripts utilised was ENST00000562631.7 (Human) and ENSGGOT00000008484.3 (Gorilla), respectively for Seq1 and Seq2.

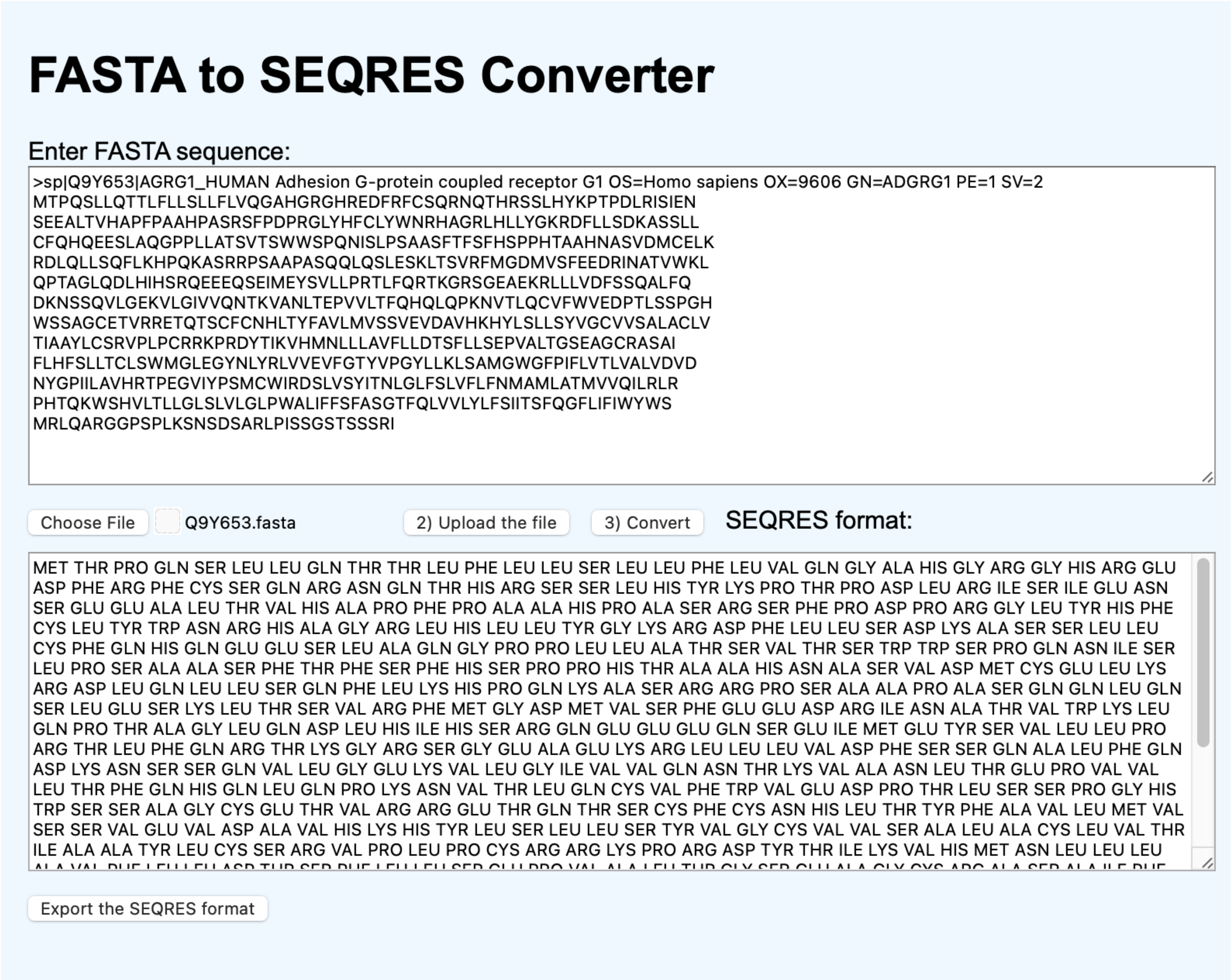


**Figure S5. FASTA to SEQRES converter on the website.**

```
>results for 693 residue sequence "sp|Q9Y653|AGRG1_HUMAN Adhesion G-protein coupled recept
MetThrProGlnSerLeuLeuGlnThrThrLeuPheLeuLeuSerLeuLeuPheLeuVal
GlnGlyAlaHisGlyArgGlyHisArgGluAspPheArgPheCysSerGlnArgAsnGln
ThrHisArgSerSerLeuHisTyrLysProThrProAspLeuArgIleSerIleGluAsn
SerGluGluAlaLeuThrValHisAlaProPheProAlaAlaHisProAlaSerArgSer
PheProAspProArgGlyLeuTyrHisPheCysLeuTyrTrpAsnArgHisAlaGlyArg
LeuHisLeuLeuTyrGlyLysArgAspPheLeuLeuSerAspLysAlaSerSerLeuLeu
CysPheGlnHisGlnGluGluSerLeuAlaGlnGlyProProLeuLeuAlaThrSerVal
ThrSerTrpTrpSerProGlnAsnIleSerLeuProSerAlaAlaSerPheThrPheSer
PheHisSerProProHisThrAlaAlaHisAsnAlaSerValAspMetCysGluLeuLys
ArgAspLeuGlnLeuLeuSerGlnPheLeuLysHisProGlnLysAlaSerArgArgPro
SerAlaAlaProAlaSerGlnGlnLeuGlnSerLeuGluSerLysLeuThrSerValArg
PheMetGlyAspMetValSerPheGluGluAspArgIleAsnAlaThrValTrpLysLeu
GlnProThrAlaGlyLeuGlnAspLeuHisIleHisSerArgGlnGluGluGluGlnSer
GluIleMetGluTyrSerValLeuLeuProArgThrLeuPheGlnArgThrLysGlyArg
SerGlyGluAlaGluLysArgLeuLeuLeuValAspPheSerSerGlnAlaLeuPheGln
AspLysAsnSerSerGlnValLeuGlyGluLysValLeuGlyIleValValGlnAsnThr
LysValAlaAsnLeuThrGluProValValLeuThrPheGlnHisGlnLeuGlnProLys
AsnValThrLeuGlnCysValPheTrpValGluAspProThrLeuSerSerProGlyHis
TrpSerSerAlaGlyCysGluThrValArgArgGluThrGlnThrSerCysPheCysAsn
```

**Figure S6. Output from another website tool for sequence conversion (www.bioinformatics.org/sms2/one_to_three).**

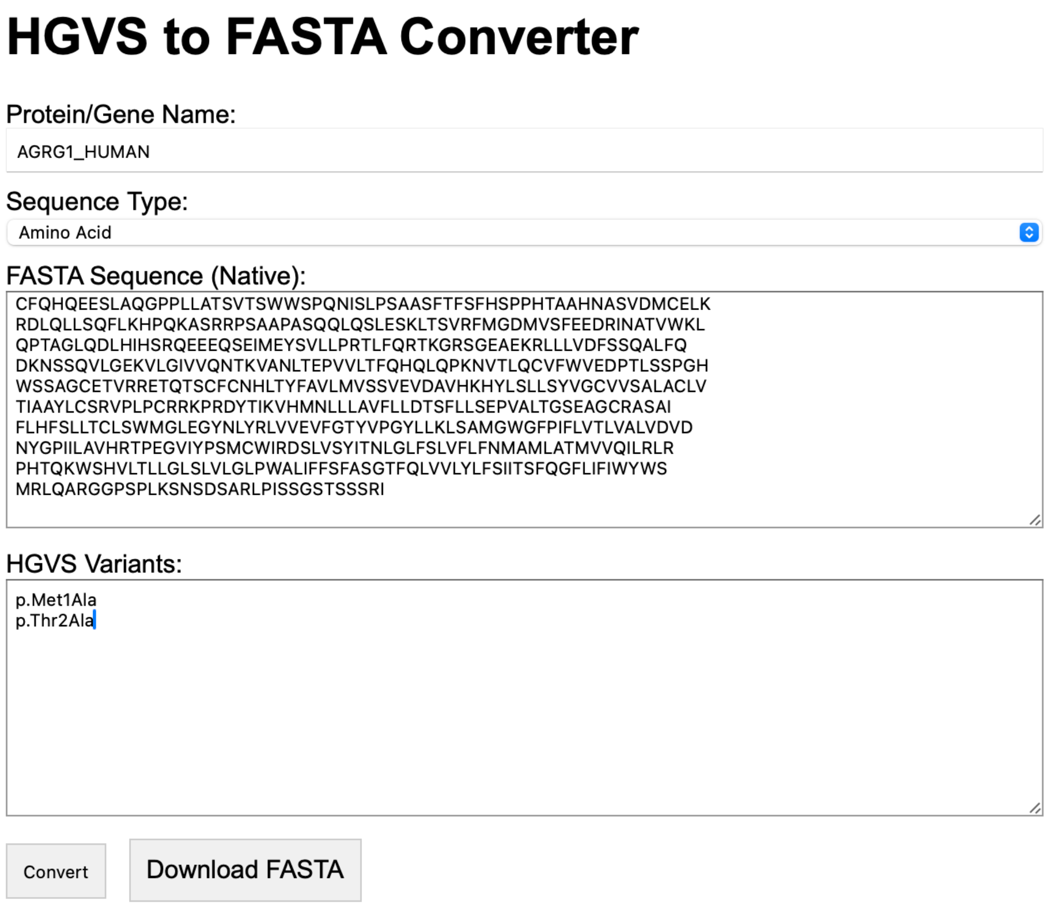


**Figure S7. User interface of the HGSV to FASTA converter for AGRG1_HUMAN protein.**

```
>AGRG1_HUMAN
MTPQSLLQTTLFLLSLLFLVQGAHGRGHREDFRFCSQRNQTHRSSLHYKPTPDLRISIEN
SEEALTVHAPFPAAHPASRSFPDPRGLYHFCLYWNRHAGRLHLLYGKRDFLLSDKASSLL
CFQHQEESLAQGPPLLATSVTSWWSPQNISLPSAASFTFSFHSPPHTAAHNASVDMCELK
RDLQLLSQFLKHPQKASRRPSAAPASQQLQSLESKLTSVRFMGDMVSFEEDRINATVWKL
QPTAGLQDLHIHSRQEEEQSEIMEYSVLLPRTLFQRTKGRSGEAEKRLLLVDFSSQALFQ
DKNSSQVLGEKVLGIVVQNTKVANLTEPVVLTFQHQLQPKNVTLQCVFWVEDPTLSSPGH
WSSAGCETVRRETQTSCFCNHLTYFAVLMVSSVEVDAVHKHYLSLLSYVGCVVSALACLV
TIAAYLCSRVPLPCRRKPRDYTIKVHMNLLLAVFLLDTSFLLSEPVALTGSEAGCRASAI
FLHFSLLTCLSWMGLEGYNLYRLVVEVFGTYVPGYLLKLSAMGWGFPIFLVTLVALVDVD
NYGPIILAVHRTPEGVIYPSMCWIRDSLVSYITNLGLFSLVFLFNMAMLATMVVQILRLR
PHTQKWSHVLTLLGLSLVLGLPWALIFFSFASGTFQLVVLYLFSIITSFQGFLIFIWYWS
MRLQARGGPSPLKSNSDSARLPISSGSTSSSRI
>AGRG1_HUMAN_variant_1
ATPQSLLQTTLFLLSLLFLVQGAHGRGHREDFRFCSQRNQTHRSSLHYKPTPDLRISIEN
SEEALTVHAPFPAAHPASRSFPDPRGLYHFCLYWNRHAGRLHLLYGKRDFLLSDKASSLL
CFQHQEESLAQGPPLLATSVTSWWSPQNISLPSAASFTFSFHSPPHTAAHNASVDMCELK
RDLQLLSQFLKHPQKASRRPSAAPASQQLQSLESKLTSVRFMGDMVSFEEDRINATVWKL
QPTAGLQDLHIHSRQEEEQSEIMEYSVLLPRTLFQRTKGRSGEAEKRLLLVDFSSQALFQ
DKNSSQVLGEKVLGIVVQNTKVANLTEPVVLTFQHQLQPKNVTLQCVFWVEDPTLSSPGH
WSSAGCETVRRETQTSCFCNHLTYFAVLMVSSVEVDAVHKHYLSLLSYVGCVVSALACLV
TIAAYLCSRVPLPCRRKPRDYTIKVHMNLLLAVFLLDTSFLLSEPVALTGSEAGCRASAI
FLHFSLLTCLSWMGLEGYNLYRLVVEVFGTYVPGYLLKLSAMGWGFPIFLVTLVALVDVD
NYGPIILAVHRTPEGVIYPSMCWIRDSLVSYITNLGLFSLVFLFNMAMLATMVVQILRLR
PHTQKWSHVLTLLGLSLVLGLPWALIFFSFASGTFQLVVLYLFSIITSFQGFLIFIWYWS
MRLQARGGPSPLKSNSDSARLPISSGSTSSSRI
>AGRG1_HUMAN_variant_2
MAPQSLLQTTLFLLSLLFLVQGAHGRGHREDFRFCSQRNQTHRSSLHYKPTPDLRISIEN
SEEALTVHAPFPAAHPASRSFPDPRGLYHFCLYWNRHAGRLHLLYGKRDFLLSDKASSLL
CFQHQEESLAQGPPLLATSVTSWWSPQNISLPSAASFTFSFHSPPHTAAHNASVDMCELK
RDLQLLSQFLKHPQKASRRPSAAPASQQLQSLESKLTSVRFMGDMVSFEEDRINATVWKL
QPTAGLQDLHIHSRQEEEQSEIMEYSVLLPRTLFQRTKGRSGEAEKRLLLVDFSSQALFQ
DKNSSQVLGEKVLGIVVQNTKVANLTEPVVLTFQHQLQPKNVTLQCVFWVEDPTLSSPGH
WSSAGCETVRRETQTSCFCNHLTYFAVLMVSSVEVDAVHKHYLSLLSYVGCVVSALACLV
TIAAYLCSRVPLPCRRKPRDYTIKVHMNLLLAVFLLDTSFLLSEPVALTGSEAGCRASAI
FLHFSLLTCLSWMGLEGYNLYRLVVEVFGTYVPGYLLKLSAMGWGFPIFLVTLVALVDVD
NYGPIILAVHRTPEGVIYPSMCWIRDSLVSYITNLGLFSLVFLFNMAMLATMVVQILRLR
PHTQKWSHVLTLLGLSLVLGLPWALIFFSFASGTFQLVVLYLFSIITSFQGFLIFIWYWS
MRLQARGGPSPLKSNSDSARLPISSGSTSSSRI
```

**Figure S8. FASTA formats for p.Met1Ala and p.Thr2Ala variants of AGRG1.** Generated by HGSV to FASTA converter on the toolkit website.

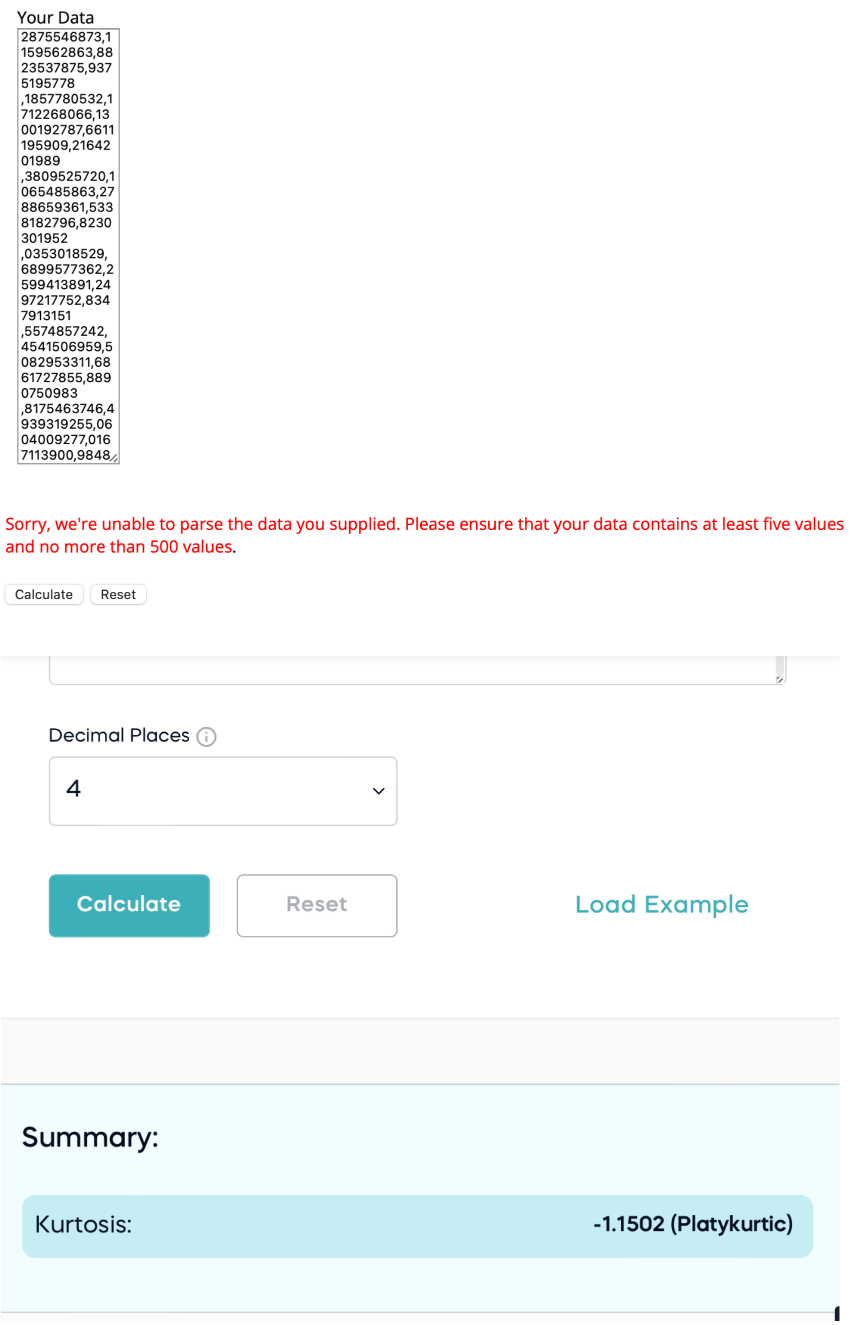


**Figure S9. Two different tools for kurtosis coefficient calculation.**
Our kurtosis calculator handles more than 500 data values compared to (www.socscistatistics.com/tests/skewness/). Below is a website that could calculate from more values but without a graphical output (exploringfinance.com/kurtosis-calculator/).

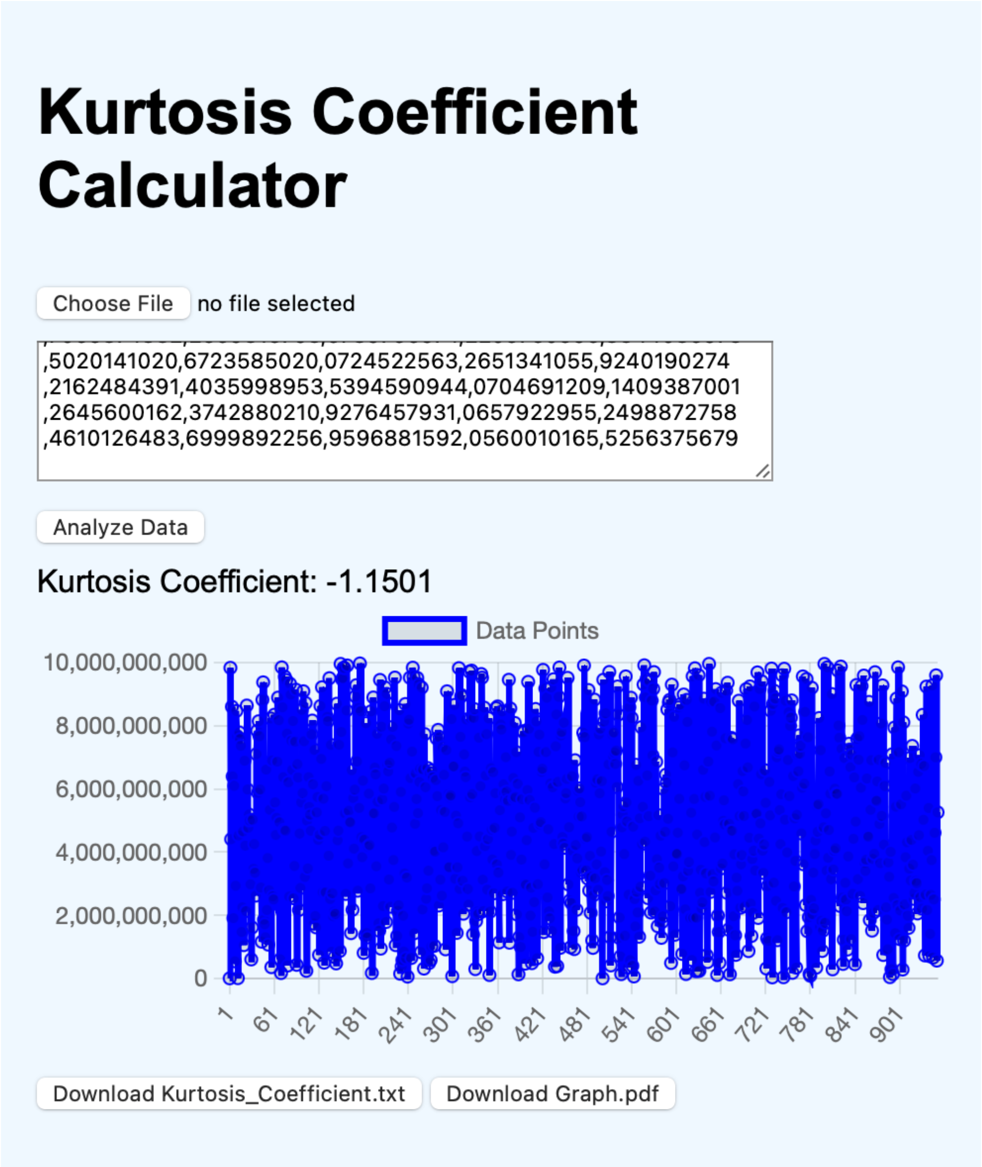


**Figure S10. Kurtosis coefficient calculator on the toolkit website.**

Kurtosis Coefficient: -1.1501

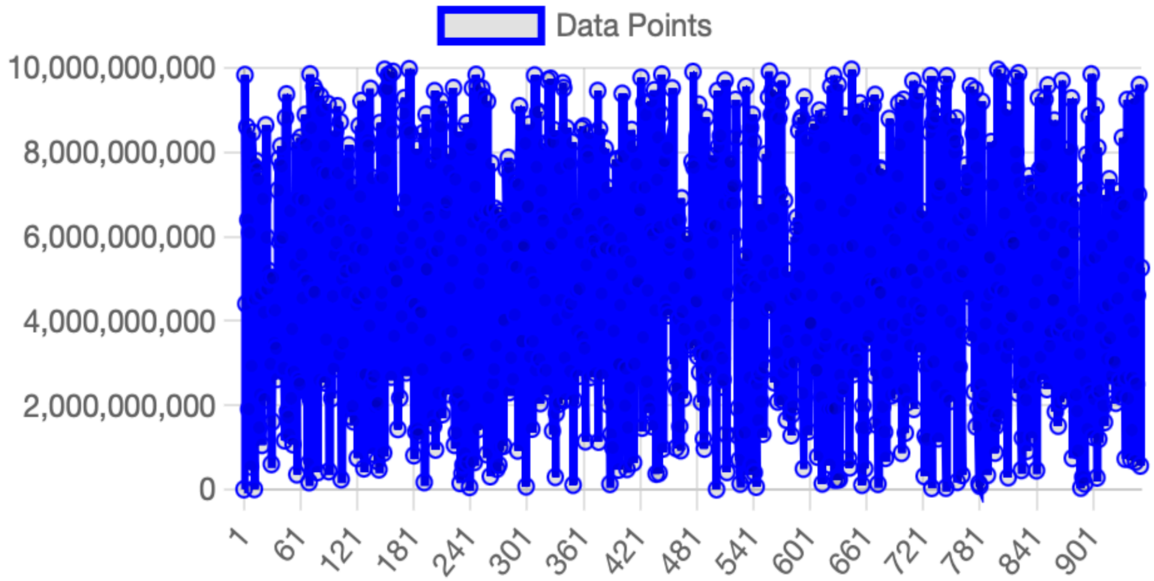


**Figure S11. Output of the kurtosis coefficient calculator using segments of Pi digits ranging from 500 to 10,000, with each segment containing 10 digits.**

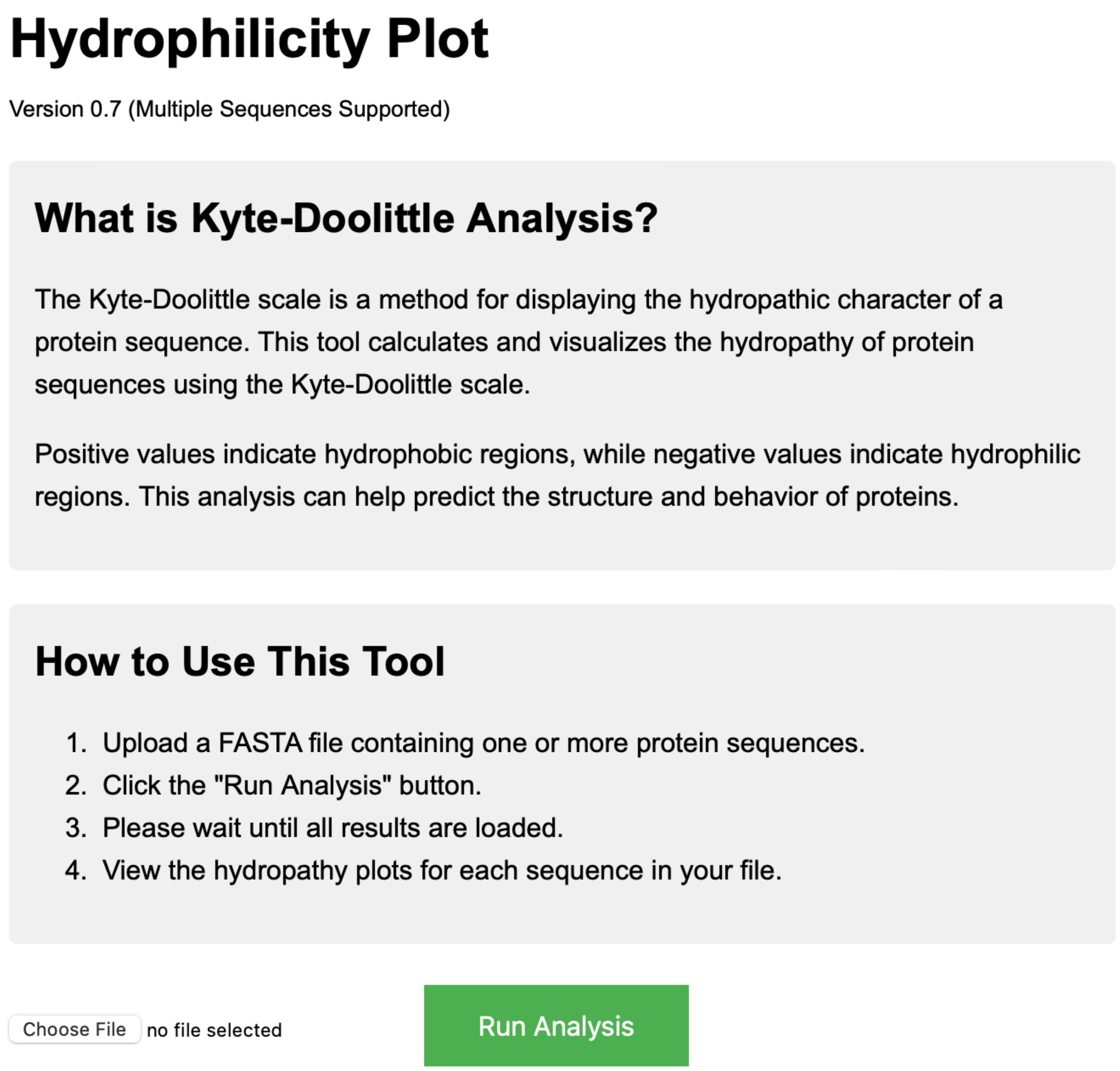


**Figure S12. The graphical user interface (GUI) for the Hydrophilicity Plot tool.** This tool is based on the Kyte-Doolittle analysis, which measures the hydropathy (hydrophobic or hydrophilic regions) of protein sequences.

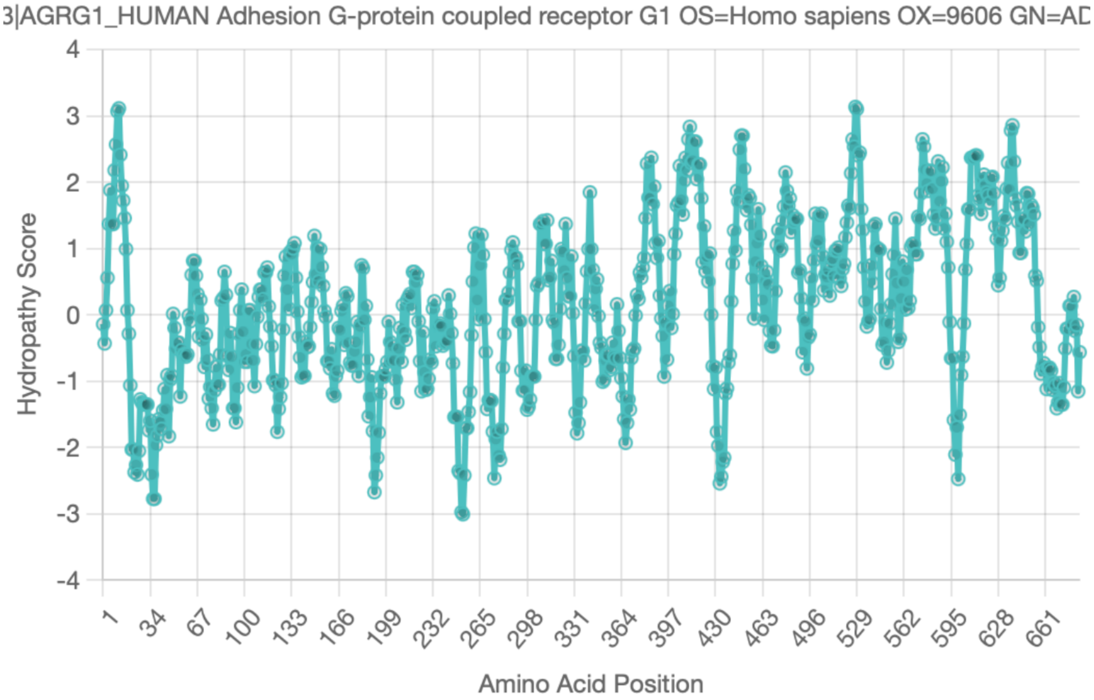


Kyte-Doolittle hydropathy plot for the sequence "sp|Q9Y653|AGRG1_HUMAN Adhesion G-protein coupled receptor G1 OS=Homo sapiens OX=9606 GN=ADGRG1 PE=1 SV=2". Positive values indicate hydrophobic regions, while negative values indicate hydrophilic regions.

**Figure S13. Output of the Kyte-Doolittle Analysis tool using AGRG1 sequence.** The numbers along the x axis represent the position of each amino acid in the protein sequence. Y-axis indicates hydropathy scores which ranges from approximately -4 to +4. Below the plot, an explation for Kyte-Doolittle values locates.
